# Herbicidal interference: glyphosate drives both the ecology and evolution of plant-herbivore interactions

**DOI:** 10.1101/2024.06.05.597659

**Authors:** Grace M. Zhang, Regina S. Baucom

**Affiliations:** Ecology and Evolutionary Biology Department, 4034 Biological Sciences Building, University of Michigan, Ann Arbor, MI 48109

**Keywords:** eco-evolutionary dynamics, herbicide resistance, herbivory resistance, plant-herbivore interactions, glyphosate

## Abstract

The coevolution of plants and their insect herbivores reflects eco-evolutionary dynamics at work— ecological interactions influence adaptive traits, which feed back to shape the broader ecological community. However, novel anthropogenic stressors like herbicide, which are strong selective agents, can disrupt these dynamics. Little is known about how the evolution of herbicide resistance may impact plant-herbivore interactions. We performed a common garden field experiment using *Ipomoea purpurea* (common morning glory) and the herbicide glyphosate (Roundup) to investigate the ecological effects of herbicide exposure on insect herbivory patterns and assess the potential evolutionary consequences. We find that plants treated with glyphosate experienced higher levels of herbivory and altered chewing herbivory damage patterns. Additionally, we found that glyphosate resistance is positively associated with herbivory resistance, and uncovered positive selection for increased glyphosate resistance, suggesting that selection for increased glyphosate resistance has the potential to lead to increased herbivory resistance. Positive selection for glyphosate resistance, coupled with the detection of genetic variation for this trait, suggests there is potential for glyphosate resistance—and herbivory resistance *via* hitchhiking— to further evolve. Our results show that herbicides can not just influence, but potentially drive the eco-evolutionary dynamics of plant-herbivore interactions.

## INTRODUCTION

Plants have coexisted and interacted with insect herbivores for more than 400 million years (Ehrlich & Raven, 1964; Labandeira & Currano, 2013), with such interactions influencing the evolution of plant traits, population dynamics, and even community stability (Myers & Sarfraz, 2017; De-la-Cruz *et al*., 2020; Agrawal & Maron, 2022). Indeed, interactions between plants and herbivores can reciprocally influence their respective evolutionary trajectories—herbivores exert selective pressure on plants, leading to the evolution and diversification of anti-herbivory defenses; in turn, evolutionary responses to herbivory can alter the ecological conditions in which plants and herbivores interact (Agrawal *et al*., 2006). Such eco-evolutionary feedback loops can occur in the context of human-mediated forms of stress like those associated with agricultural regimes, urbanization, and climate change (Turcotte *et al*., 2017; Hamann *et al*., 2021; Santangelo *et al*., 2022). However, how these human-mediated stressors alter plant-herbivore interactions and thus the broader eco-evolutionary dynamic between plants and their insect herbivores remains a significant gap in our knowledge.

Chemical herbicides are a relatively novel form of human-mediated selection used in agricultural ecosystems to control and eradicate weedy plants (Shaner, 2014). Unfortunately, plants have quickly adapted to herbicide use; to date there are hundreds of weed species that are considered either resistant or tolerant to some form of herbicide (Baucom & Mauricio, 2004; Vila-Aiub *et al*., 2009; Délye *et al*., 2013). Most work addressing natural weeds and herbicide use examines some aspect of the evolution of resistance or tolerance in plant populations—whether focusing on the frequency of resistance or tolerance, the genetic basis of either defense trait, or the potential for fitness costs associated with herbicide defense (Baucom, 2019). Far fewer studies consider the consequences of herbicides on non-target organisms such as herbivorous insects or pollinators within the weed community (Motta *et al*., 2018, 2020). While some research has examined how herbicide exposure may directly influence insect development, fitness, and immune responses (Schneider *et al*., 2009; Capinera, 2018; Baglan *et al*., 2018; Smith *et al*., 2021), the potential for indirect effects of herbicide—where herbicide exposure alters or changes some aspect of the plants that subsequently impacts insects or other community members (Fuchs *et al*., 2021)—is less understood. This is a notable knowledge gap because plant chemistry, physiology, and phenology are directly altered by herbicide exposure (Baucom *et al*., 2008; Londo *et al*., 2014; Freitas-Silva *et al*., 2022); additionally, aspects of plant size and the mating system have evolved along with herbicide resistance (Van Etten *et al*., 2016; Kuester *et al*., 2017). Each of these changes— whether trait changes due to exposure, or trait alterations due to correlated evolution with herbicide resistance—has the potential to interfere with established interactions between plants and insect herbivores, pollinators, and other organisms.

There is evidence that plant herbivory specifically may be impacted by herbicide. For example, in glyphosate-tolerant *Beta vulgaris* (sugar beet), glyphosate application led to an increase in the herbivore *Myzus persicae* (green peach aphid) (Dewar *et al*., 2000). Similarly, *Abutilon theophrasti* (velvetleaf) exposed to low doses of the herbicide dicamba show an elevated abundance of the phloem-feeding *Bemisia tabaci* (silverleaf whitefly) (Johnson & Baucom, 2022). The consequences of such increased herbivory experienced by herbicide-exposed plants are generally unknown on both the part of the plant and the insect herbivore. For example, does the increased abundance of an herbivore due to herbicide application lead to greater selective pressure than the plant population would normally experience? Or do different types of feeding result, potentially influencing both the trajectory of herbivory resistance in plants and the herbivore population/community in unexpected ways? It is also unknown if plants exposed to herbivory and herbicide simultaneously may eventually evolve higher levels of resistance to both stressors.

At the same time, there is also the possibility that plants may experience a trade-off between defense to herbicide and herbivory, which could lead to a constraint on the evolution of increased levels of either trait (Simms & Rausher, 1987). Indeed, there have been trade-offs detected between constitutive and induced anti-herbivory defenses (Koricheva *et al*., 2004), between resistance to different herbivore species (Agrawal *et al*., 1999), and between the defense strategies of resistance and tolerance to the same threat (van der Meijden *et al*., 1988; Fineblum & Rausher, 1995; Baucom & Mauricio, 2008). However, the evidence that trade-offs exist between herbivory and herbicide defense is more limited; one example showed that *Amaranthus hybridus* (smooth pigweed) plants resistant to the herbicide triazine were more susceptible to the specialist herbivore *Disonycha glabrata* (striped flea beetle) as compared to triazine-susceptible plants (Gassmann & Futuyma, 2005). It is possible that trade-offs between herbicide and herbivory resistance exist but remain undetected only due to the small number of studies on this topic to date.

Here, we consider plant-herbivore interactions in the context of an herbicide-exposed plant population. As human activity has become a primary driver of global change, human-mediated stressors may impact plant-herbivore interactions, thereby potentially altering plant evolutionary responses, which could subsequently feed back to and reshape the ecological context within which plant herbivory is occurring.

Our goal is to investigate the effects of a novel but strong selective agent—herbicide—on long-standing plant-herbivore relationships using *Ipomoea purpurea* (common morning glory). We specifically examine both the ecological effects of herbicide exposure on insect herbivory and the potential for evolutionary consequences of herbicide resistance on the evolution of herbivory resistance. To this end, we ask the following questions regarding ecological impacts: 1) Does glyphosate application affect insect herbivory levels and/or chewing damage patterns? If so, glyphosate has the potential to alter plant-herbivore interactions and thus indirectly moderate the insect herbivore community by favoring the feeding of certain herbivores over others. We further ask: 2) Is there a trade-off between glyphosate and herbivory resistance? The existence of a trade-off would indicate that the two forms of resistance may be mutually exclusive, which may evolutionarily constrain the evolution of either form of resistance. Finally, we examine the potential evolutionary consequences and ask: 3) Is there potential for these resistance traits to further evolve? If so, and a trade-off is present, then not only may plant populations start to diverge based on their resistance strategies, but their associated insect herbivore communities may diverge as well, as herbivores adapt to the evolving plants that form their diet. By answering these questions, our study begins to uncover the eco-evolutionary effects that herbicide may exert on plant-herbivore communities.

## MATERIALS AND METHODS

### Study system

*Ipomoea purpurea* (L.) Roth (Convolvulaceae), the common morning glory, is an annual weed commonly found in agricultural fields and disturbed roadsides of the US midwest and Southeast. Seeds germinate from early summer to mid-autumn and flowers emerge four to six weeks after germination. Daily flowering continues until the plant is killed by the first frost (Debban *et al*., 2015). The species has a mixed-mating system and is typically pollinated by bumblebees, honeybees, and syrphid flies. Fruits mature about four weeks after pollination and plants are prolific, capable of producing up to 8000 seeds a season (Chaney & Baucom, 2014). Previous work has shown that insect herbivores select for higher levels of herbivory resistance in *I. purpurea* (Simms & Rausher, 1989; Tiffin & Rausher, 1999), and that this species has naturally evolved higher levels of resistance to the herbicide glyphosate (Baucom & Mauricio, 2008; Kuester *et al*., 2015).

### Field experiment

On June 3, 2021, replicate *Ipomoea purpurea* seeds were planted in a common garden at the Matthaei Botanical Gardens in Ann Arbor, Michigan. Four to eight replicate seeds from 101 maternal lines representing 18 natural populations sampled from the US Southeast and Midwest in 2012 were selected to form a synthetic population of 1612 individuals that exhibited a wide range of resistance to glyphosate (Schluter, 1988; Kuester *et al*., 2015). Seeds were planted in a randomized block design with two blocks containing both a control (non-herbicide) and treatment (herbicide) environment. Four weeks after germination, we conducted a leaf count of all plants as a proxy for plant size. We then applied glyphosate at a rate of 1.5 kg ai/ha to the treatment group six weeks after germination, which is higher than the typical field dose (1.12 kg ai/ha (Baucom & Mauricio, 2004)). Thus, in this experiment we are using a rate of glyphosate that should differentiate resistant from susceptible lineages. To score glyphosate damage, which is typified by a yellowing and browning of the leaves, the percentage of damaged leaves was estimated for every treated plant two weeks after glyphosate application as the number of damaged leaves divided by the total number of leaves (following Baucom and Mauricio 2008). Four weeks after treatment, we scored every treated plant as either dead or alive as a binomial response (dead = 0, alive = 1).

Due to vole damage as well as death following glyphosate exposure in the treatment group, the number of plants in both the control and treatment groups were reduced by the time of seed collection. In the control treatment, 437 plants survived to produce seed, and 306 plants in the glyphosate treatment survived to produce seed. We were able to collect herbivory damage on 439 and 307 plants in the control and glyphosate treatments, respectively (see below for details). Our estimate of fitness in this experiment is survival to seed production (no seeds = 0, at least one seed = 1). Preliminary analyses indicated there was no evidence of maternal line variation for vole damage (maternal line effect: 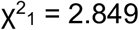, p = 0.091) and as such we did not examine vole damage further.

To assess insect herbivory damage, three leaves per plant were photographed a month after glyphosate application and analyzed using the app LeafByte (Getman-Pickering *et al*., 2020). The percentage of leaf area consumed was recorded. We elected to take a conservative measure of herbivory since herbivory and glyphosate damage were at times overlapping and difficult to delineate, especially on the leaf margins (Fig. S1). Specifically, when extracting herbivory data from photos of glyphosate-treated plants, we excluded areas of the leaf in which we could not confidently differentiate herbivory from glyphosate damage. We note that although our conservative approach could potentially lead to reduced estimates of herbivory, it would not influence our measure of herbicide resistance (*i*.*e*., 1 - % of leaves showing damage across the entire plant; Fig. S1).

To characterize the type of herbivory damage found on the photographed leaves, we referred to the *Guide to Insect (and Other) Damage Types on Compressed Insect Fossils. Version 3*.*0*. (Labandeira *et al*., 2007). Each leaf was scored for the presence/absence of three categories of chewing damage as defined in the guide: hole-feeding, margin-feeding, and surface-feeding (Fig. 1). We then calculated the proportion of leaves of each plant that showed the presence of each of the three categories of herbivory damage to statistically compare the prevalence of damage types between the control and treatment groups.

**Figure 1.**
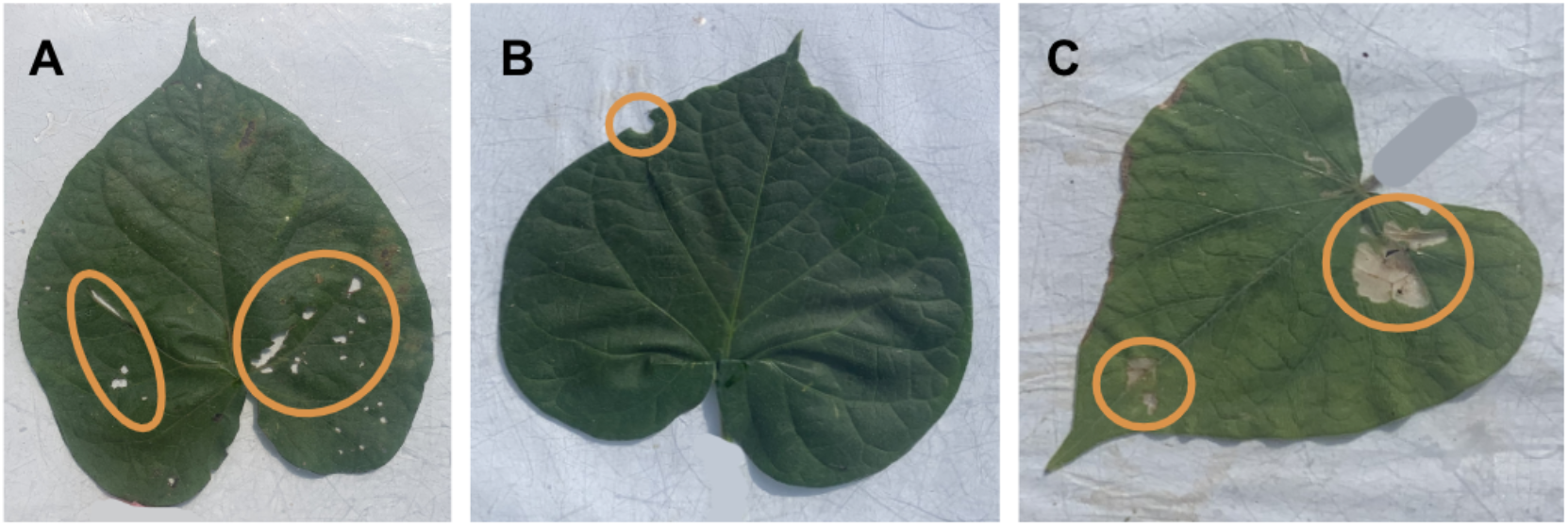
Instances of herbivory damage, circled in orange, were categorized into three broad categories: (A) hole-, (B) margin-, and (C) surface-feeding.

### Statistical analysis

#### Effect of glyphosate treatment on herbivory and damage types

Resistance was operationally defined as 1 - *p*, where *p* is the amount of damage. For glyphosate resistance, *p* was the percentage of glyphosate-damaged leaves per plant (Baucom & Mauricio, 2008), while for herbivore resistance, *p* was the average percentage of leaf area consumed per plant (Rausher & Simms, 1989).

We used the program R (version 4.2.1) (R Core Team, 2023) to conduct all statistical analyses. Variables were first transformed using the appropriate method as identified by the *bestNormalize* package (Peterson, 2021). We first used the *lme4* package (Bates *et al*., 2015) to construct a linear mixed model for herbivory resistance. We included both control and treated plants and used the following model to test for a maternal line effect (and thus the presence of genetic variation) and a treatment effect:

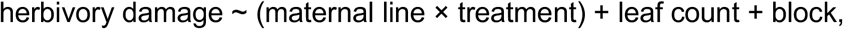

with block, maternal line, and the interaction between maternal line and treatment block as random effects. To test the fixed effects, we used the Satterthwaite approximation to determine the degrees of freedom. To assess the significance of the random effects, we removed each random effect and compared the simplified model to the full model using a χ^2^ test (df = 1).

Next, we determined if glyphosate treatment could result in herbivore community-level changes by examining the effects of glyphosate application on the type of chewing damage. The damage type categories of hole-, margin-, and surface-feeding were log-transformed prior to analysis. We then assessed if there was an effect of glyphosate application on the frequency of damage type occurrences by fitting linear mixed models with the damage type frequency of a particular category as the response variable and treatment as a fixed effect, while maternal line, the maternal line by treatment interaction, and block were included as random effects. As above, we compared the full to simplified models using a χ^2^ test (df = 1) to assess the significance of the random effects in the model.

#### Relationship between glyphosate resistance and herbivory

We tested for a relationship between glyphosate and herbivory resistance by constructing a linear mixed model using glyphosate-treated plants with the following model:

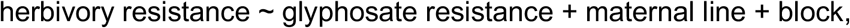

with maternal line and block as random effects. We also calculated Pearson’s correlation between the maternal line averages of glyphosate and herbivory resistance to determine if there was a genotypic correlation between the two measures of resistance.

Similarly, we determined whether there was a relationship between glyphosate resistance and resistance to each of the three damage type categories by constructing linear mixed models with the frequency of each damage type occurrence as the response variable. Glyphosate resistance was tested as a fixed effect, and maternal line and block were tested as random effects. Additionally, we calculated genotypic Pearson’s correlations between the maternal line average of glyphosate resistance and resistance to each of the three damage type categories.

Next, we assessed the potential for genetic variation underlying glyphosate resistance in two ways. First, we constructed a linear mixed model with glyphosate resistance as the response variable:

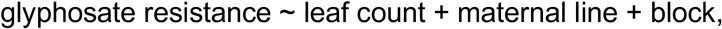

with leaf count (as a plant size covariate) as a fixed effect, and maternal line and block as random effects. Second, we constructed a generalized linear mixed model using the binomial link family with plant death as the response variable (*i*.*e*., either dead (0) or alive (1) after glyphosate treatment):

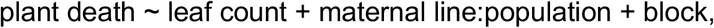

with leaf count as a fixed effect, and maternal line nested in population and block as random effects. To assess the effects of maternal line in both models, we constructed reduced models that did not include maternal line, and used a χ^2^ test (df = 1) to compare models.

#### Direct and indirect effects of glyphosate on seed set

We next assessed the relative influence of glyphosate and herbivory resistance on fitness to determine which resistance trait might be more likely to evolve in this species, and determined the direct and indirect relationships between glyphosate treatment, glyphosate resistance, herbivory resistance, and fitness in the form of seed set. To do so we constructed a structural equation model (SEM) with data from both treatment environments using the *piecewiseSEM* package in R (Lefcheck, 2016), including treatment, glyphosate resistance, herbivory resistance, and seed set as variables. Maternal line and block were included as random effects. To assess the model fit, we calculated the Fisher’s *C* statistic and compared it to a χ^2^ distribution. Specifically, we used mixed models to assess the following relationships:

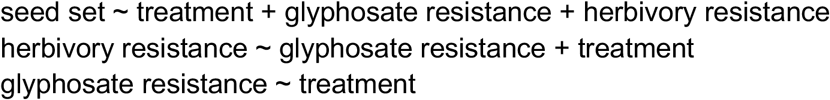

We first built an SEM that included relationships between all four variables (*i*.*e*., treatment, herbivory resistance, glyphosate resistance, seed set). This model, Model A, resulted in a saturated SEM that did not allow us to assess model fit. Thus, we constructed an SEM that omitted the relationship between herbivory resistance and seed set (Fig. 4), Model B, which best fitted our data. We compared the AIC scores of Model A and Model B as part of model selection and found that Model B had the lower AIC score. Accordingly, we proceeded with Model B as our hypothesized SEM.

We specifically examined the effects of glyphosate resistance on herbivory resistance rather than exploring a correlational relationship because we were interested in how resistance to a new stressor (*i*.*e*., glyphosate) may alter resistance to a preexisting, long-standing stressor (*i*.*e*., insect herbivory). Moreover, by separately examining how glyphosate *application* (*i*.*e*., treatment) and glyphosate *resistance* may impact herbivory resistance, we can parse treatment effects from the direct effects of glyphosate resistance and evaluate the relative strength and direction of those relationships independently. To be thorough, we tested an SEM modified from Model B to include the correlational relationship between herbivory resistance and glyphosate resistance. This model (Model C) had a higher AIC score than Model B, further validating Model B as the best-fitting SEM.

## RESULTS

### Glyphosate alters the amount of herbivory and the pattern of damage

We first examined the potential that plants treated with glyphosate showed altered rates of herbivory. We found that glyphosate exposure led to 112.7% more herbivory damage compared to control plants (F_1_ = 144.26, p < 0.001; Fig. 2a; Table S1). We note that this estimate of herbivory damage is potentially conservative since we excluded ambiguously damaged areas from the estimate of chewing leaf herbivory. We further assessed the ambiguous damage in two ways: first, we determined if the excluded regions were more likely glyphosate or herbivory damage, and second, we examined herbivory damage as a qualitative score rather than an amount, as a way to verify that glyphosate-treated plants experience more chewing herbivory. We found that plants that were more susceptible to glyphosate had more leaves with ambiguous damage (F_1_ = 35.113, p < 0.001), and likewise found that there was no relationship between herbivory damage and the estimated ambiguous damage (F_1_ = 1, p = 0.457; Fig. S1). This suggests that the excluded areas are likely glyphosate rather than herbivory damage, and that our conservative estimate of herbivory is appropriate. Furthermore, when we compared treated and control plants using the presence/absence of herbivory damage as our metric, we found that leaves of treated plants were 21% more likely to show herbivory than control plants (see Fig. S1 legend for details of analyses), further supporting our result of increased rates of herbivory on plants exposed to glyphosate.

**Figure 2.**
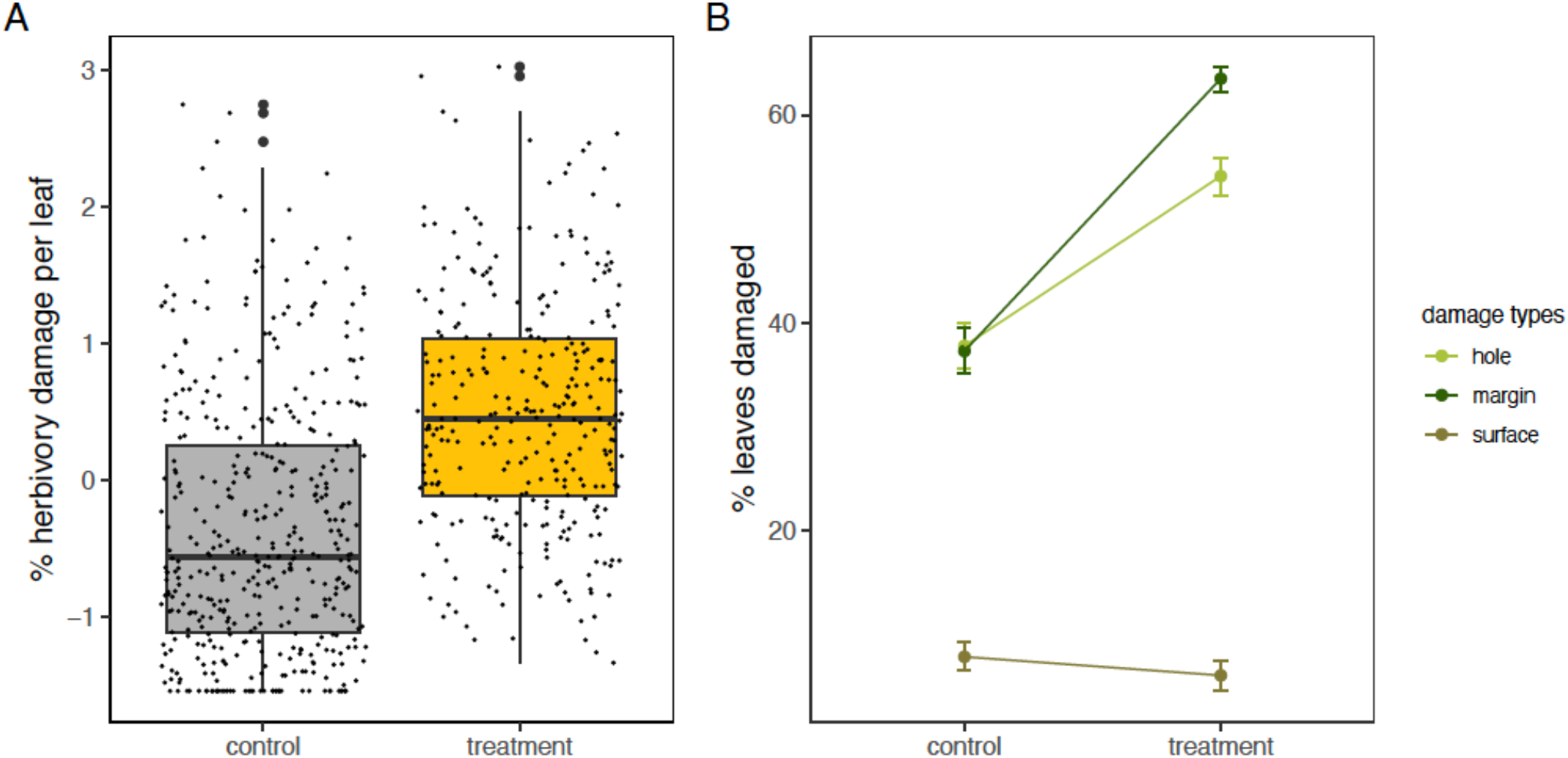
(A) Glyphosate-treated plants experience higher levels of insect herbivory than control plants (F_1_ = 131.950, p < 0.001). Each point is an individual plant (treated plants N = 307, control plants N = 439). (B) Glyphosate-treated plants experience more occurrences of hole-feeding (F_1_ = 100.490, p < 0.001) and margin-feeding (F_1_ = 346, p < 0.001). Instances of surface-feeding is lower in treated plants as compared to control plants (F_1_ = 5.208, p < 0.023).

When examining herbivory damage type categories (hole-, margin-, and surface-feeding), we found that glyphosate-treated plants showed a higher proportion of leaves with hole- and margin-feeding (hole feeding, glyphosate vs. control: 74.1% vs. 50.0%, F_1_ = 100.49, p < 0.001; margin feeding: 89.8% vs. 48.9%, F_1_ = 346.00, p < 0.001), but a lower proportion of leaves showed surface-feeding compared to control plants (glyphosate vs. control: 7.2% vs. 9.4%, F_1_ = 5.21, p = 0.023, Table S2; Fig. 2b).

We did not find evidence of a maternal-line effect for the amount of herbivory damage (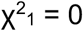, p = 1, Table S1), meaning that we find no evidence of genetic variation for herbivory resistance in this study population. However, we found a significant interaction between maternal line and treatment (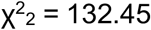, p < 0.001), suggesting that maternal lines exposed to glyphosate showed different levels of herbivory damage (*i*.*e*., there was a genotype by treatment interaction on the amount of herbivory damage). We did not detect genetic variation for the three herbivory damage type categories of hole-, margin-, and surface-feeding, but we did find a significant interaction between maternal line and treatment for hole- and margin-feeding (hole: 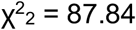, p < 0.001; margin: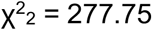, p < 0.001; Table S2).

### Relationship between glyphosate and herbivory resistance

We next assessed the potential for a trade-off between glyphosate and herbivory resistance. Instead of a negative relationship between the two defense traits, which would indicate a trade-off, we found strong evidence of a *positive* relationship between herbivory and herbicide resistance, with each trait estimated as a maternal line average (r^2^ = 0.346, p < 0.001; F_1_ = 13.94, p < 0.001; Fig. 3a). We further identified the presence of genetic variation for glyphosate resistance, both in terms of the likelihood of plant death (maternal line effect: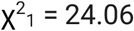, p < 0.001) and the proportion of leaves damaged due to glyphosate application (maternal line effect: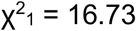, p < 0.001; Table S3).

**Figure 3.**
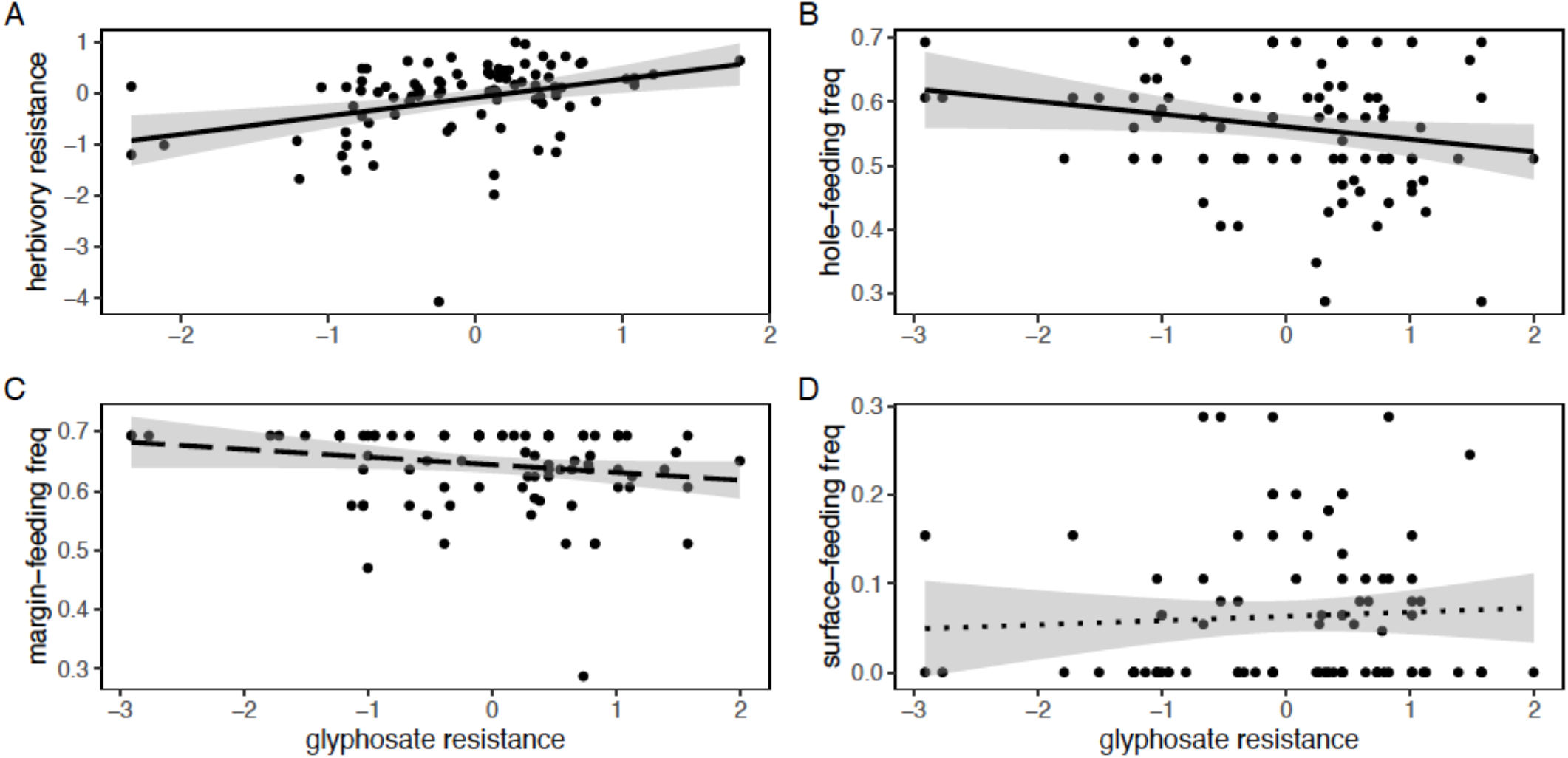
(A) Glyphosate resistance and herbivory resistance show a positive genotypic relationship (correlation r^2^ = 0.346, p < 0.001). (B) Glyphosate resistance has a negative genotypic relationship with the frequency of hole-feeding occurrences (correlation r^2^ = −0.210, p < 0.045). (C) There may be a negative genotypic relationship between glyphosate resistance and the frequency of margin-feeding occurrences (correlation r^2^ = −0.192, p < 0.068), and (D) no relationship was detected between glyphosate resistance and the frequency of surface-feeding occurrences (correlation r^2^ = 0.057, p < 0.590).

We likewise found that glyphosate resistance is negatively correlated with the frequency of hole-feeding (r^2^ = −0.210, p = 0.045; Fig. 3b) and may be negatively correlated with the frequency of margin-feeding (r^2^ = −0.190, p = 0.068; Fig. 3c). There was no relationship detected between glyphosate resistance and the frequency of surface-feeding (r^2^ = 0.057, p = 0.590; Fig. 3d).

### Direct and indirect effects of glyphosate on seed set

To determine if the resistance traits had a significant relationship with fitness, as well as examine the direct and indirect effects of glyphosate on both herbivory and the likelihood to set seed, we constructed an SEM using all experimental plants (N = 746) to test the relationships between treatment, herbivory resistance, herbicide resistance, and seed set. We compared a saturated model (Model A) that included all relationships with a simplified model that did not have a relationship between herbivory resistance and seed set (Model B). Model B provided a fit to the data (*C* = 0.47, p = 0.79, df = 2), and had a lower AIC value than Model A (4074.785 vs 4072.855, Model A vs. Model B respectively). Finally, we fit an SEM specifying correlated error between glyphosate resistance and herbivory resistance which assumes both variables are influenced by some underlying process. While this model (Model C) also fit the data (*C* = 0.47, p = 0.79, df = 2), it had a higher AIC score than Model B (4072.855 vs 4079.929, Model B vs Model C, respectively). We thus chose to proceed with the simplified model (Model B, Fig. 4, Table S4).

**Figure 4.**
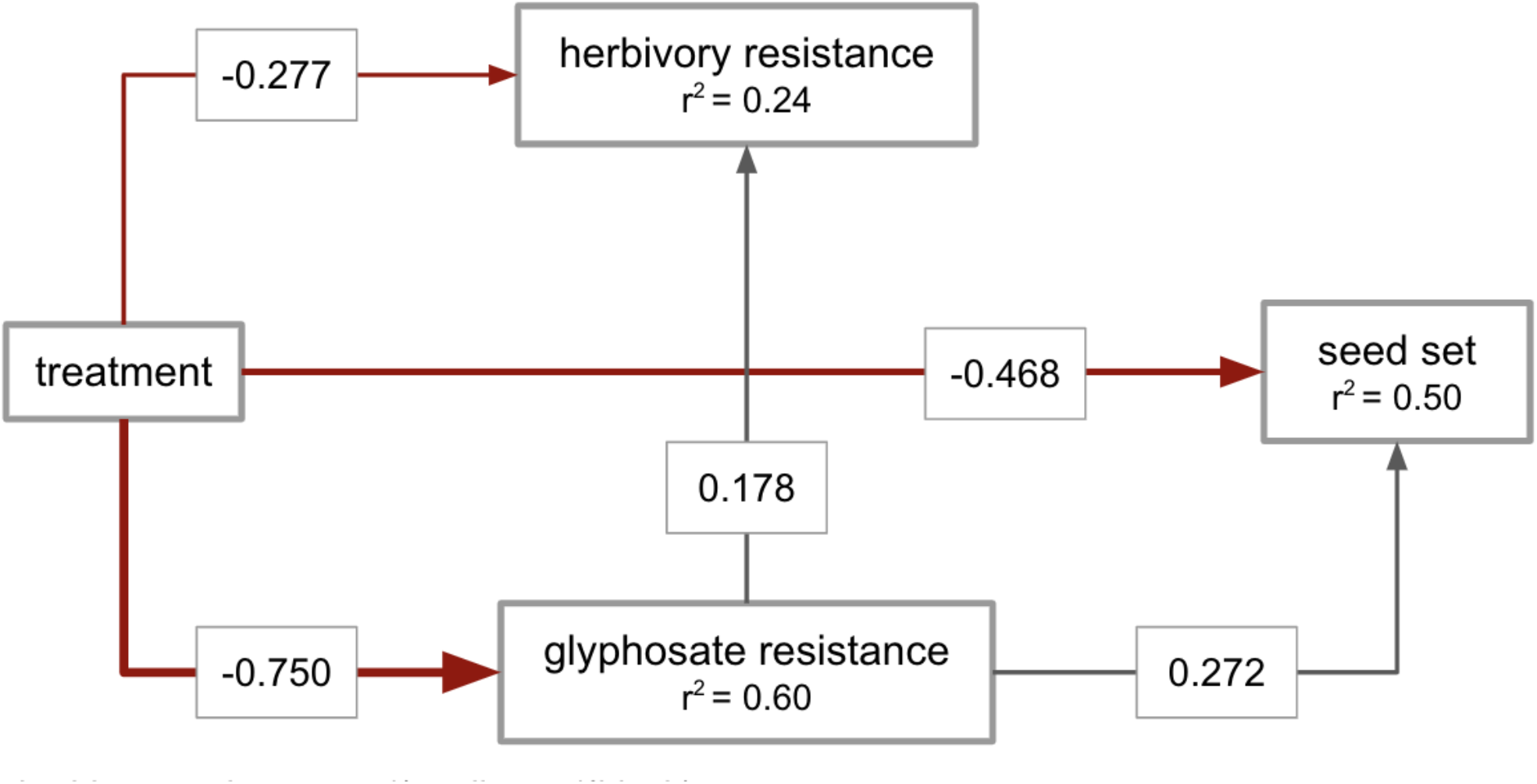
Structural equation model of the relationships between glyphosate treatment, herbivory resistance, glyphosate resistance, and seed set (Model B). Boxes represent measured variables; arrows show the unidirectional relationships between variables. Dark gray arrows show positive relationships and red arrows show negative relationships. Thickness of the arrows reflect the magnitude of the standardized β coefficients, which are shown in gray boxes along the arrows. All paths shown were highly statistically significant (p < 0.001). Conditional r^2^ values are shown within the box of each response variable. This model fit well, with a Fisher’s C score 0.47 (p = 0.79, df = 2) and an AIC score of 4072.855. Treatment has a necessarily negative effect on glyphosate resistance because resistance was calculated as 1 - *p* for the plants in both environments. In the control environment, where there was no glyphosate treatment and therefore no damage (*i*.*e*., *p* is zero), plants had a glyphosate resistance value of 1. In the treatment environment, plants experienced some level of damage, and therefore their glyphosate resistance value was less than 1. This negative treatment effect on glyphosate resistance is thus reflected in the SEM.

The selected SEM showed that glyphosate treatment was associated with lower herbivory resistance (standardized β = −0.277, df = 743, p < 0.001), lower glyphosate resistance (standardized β = −0.750, df = 710, p < 0.001), and lower seed set (standardized β = −0.468, df = 747, p < 0.001), meaning that plants exposed to glyphosate showed higher levels of herbivory (as in Fig. 2a), experienced glyphosate damage (and therefore had lower glyphosate resistance in comparison to plants in the control environment), and were less likely to set seed, respectively. We further found that glyphosate resistance was positively associated with herbivory resistance (standardized β = 0.178, df = 743, p < 0.001), as previously captured by earlier analyses (Fig. 3). Finally, we found a significant and positive relationship between glyphosate resistance and seed set (standardized β = 0.272, df = 747, p < 0.001), meaning that the greater the level of glyphosate resistance, the higher the likelihood the plant would set seed following glyphosate exposure.

It is important to note that because the hypothesized SEM with the lowest AIC score (Model B) included a direct path from glyphosate resistance to herbivory resistance (rather than a noncausal, correlational relationship between the two resistance traits, *i*.*e*., Model C), and that the standardized regression coefficient between the traits was significant and positive, our model suggests that glyphosate resistance has a direct, positive, causal effect on herbivory resistance (direct effect of glyphosate resistance on herbivory resistance, standardized β = 0.178). Further, by performing the SEM, we are able to deconstruct the influence of glyphosate on both herbivory resistance and seed set by examining the direct, indirect, and total effects of the herbicide on the traits. For herbivory resistance, we find that direct effect of glyphosate exposure explains the majority of the total effect on this trait (total effect (−0.411) = direct effect (−0.277) + indirect effect through glyphosate resistance (−0.75 × 0.178 = −0.136)). Finally, our SEM uncovered a strongly negative total effect of glyphosate on the likelihood of setting seed (−0.672) with the majority of the total effect stemming from direct rather than indirect effects (direct effect (−0.468) + indirect effect (−0.75 × 0.272 = −0.204)).

## DISCUSSION

While it has long been appreciated that interactions between plants and herbivores lead to coevolutionary dynamics between the two (Ehrlich & Raven, 1964; Bruce, 2015), we are just beginning to understand how human-mediated abiotic factors may disrupt plant-herbivore interactions and thus their evolutionary consequences (Hamann *et al*., 2021). As an anthropogenic stressor, herbicide has the potential to disrupt natural plant-herbivore interactions, leading to both ecological and evolutionary consequences. Our common garden study begins to uncover how herbicide exposure may alter established plant-herbivore dynamics. We find that glyphosate exposure leads to increased insect herbivory and altered feeding patterns, and we likewise demonstrate that glyphosate resistance is positively correlated with herbivory resistance. Coupled with our findings that there is genetic variation for glyphosate resistance, and a relationship between glyphosate resistance and fitness, it is possible that consistent selection imposed by glyphosate could lead to increased herbicide *and* herbivory resistance. Our work additionally highlights both direct and indirect effects of herbicide exposure on plant-insect interactions: herbivory levels and feeding patterns are directly altered following glyphosate exposure, whereas herbivory levels may be indirectly influenced *via* selection for increased levels of herbicide resistance. Below, we discuss the ecological effects of glyphosate exposure on herbivory, the possible mechanistic link between glyphosate resistance and herbivory resistance, and the evolutionary potential of the two forms of resistance.

### Glyphosate treatment alters herbivory patterns

Our study shows that glyphosate application increases the level of herbivory experienced by *Ipomoea purpurea*. This is in line with work showing an increased abundance of piercing-sucking herbivores on herbicide-treated plants—specifically, increased abundance of *Myzus persicae* on glyphosate-treated *Beta vulgaris*, increased *Bermisia tabaci* larvae on *Abutilon theophrasti* exposed to dicamba drift (Dewar *et al*., 2000; Johnson & Baucom, 2022), and increased *Nilaparvita lugens* on *Oryza sativa* treated with one of four herbicides (butachlor, metolachlor, bentazone, oxadiazon; Wu *et al*., 2001). Together, these results indicate that increased herbivory following herbicide exposure is a general phenomenon that is not specific to a particular herbicide or plant species. Our results expand this finding to include chewing insect herbivory.

One likely explanation for the generalized phenomenon of increased herbivory after herbicide exposure is that herbicide inhibits plant defense. The herbicide glyphosate inhibits 5-enolpyruvylshikimate-3 phosphate synthase (EPSP synthase), an important enzyme in the chorismate-producing shikimate pathway (Amrhein *et al*., 1980). Chorismate is a critical precursor molecule in the biosynthesis of essential aromatic amino acids that form the basis of many plant chemical defenses against herbivory, including volatile organic compounds (VOCs), tannins, lignins, and glucosinolates (Fuchs *et al*., 2021; Freitas-Silva *et al*., 2022). Glyphosate exposure likely alters and/or suppresses the biosynthesis of these anti-herbivory compounds, leading to increased levels of herbivory in treated plants. For example, glyphosate-susceptible soybean treated with sublethal doses of glyphosate had lower levels of lignin—an important component of physical anti-herbivory defenses (Xie *et al*., 2018)—compared to untreated plants (Marchiosi *et al*., 2009). It is likely other categories of chemical plant defense downstream of the shikimate pathway are similarly reduced after glyphosate exposure. For example, phytohormones that contribute to plant defense biosynthesis such as salicylic acid and indole-3-acetic acid are downstream of the shikimate pathway; their biosynthesis is wholly dependent on the amount of chorismate available. Additionally, salicylic acid and indole-3-acetic acid also engage in phytohormone crosstalk with other compounds critical to anti-herbivore defense, like jasmonic acid and ethylene (Fuchs *et al*., 2021). Thus, by inhibiting the shikimate pathway, glyphosate could disrupt many forms of plant herbivore defense. More work will be necessary to understand precisely how glyphosate influences herbivory at the mechanistic level.

In addition to the increased amounts of herbivory damage, we found that glyphosate-treated plants experienced altered patterns of herbivory damage, and that an increased frequency of hole- and margin-feeding in treated plants contributed to the overall increase in herbivory damage. Comparatively, surface-feeding declined on glyphosate-treated plants. This change in the pattern of feeding could be due to the disruption of plant defenses, such as VOCs (Freitas-Silva *et al*., 2022), or caused by a change to plant nutrient levels and carbon metabolic processes (Orcaray *et al*., 2012). For example, glyphosate-treated *Zea mays* (maize) showed lower levels of VOCs, making them more appealing to females of the parasitic wasp *Microplitis rufiventris* (D’Alessandro *et al*., 2006), an effect that could potentially lead to an altered herbivore community and thus different patterns of feeding.

The consequences of increased or altered herbivory given herbicide exposure are unknown, but given the change in herbivory damage patterns, we speculate that such an effect could lead to a shift in the community composition of insect herbivores, which could then lead to changes in the types of plant anti-herbivory defense traits that are under selection. Further, given that herbivores can mediate plant traits that are not directly associated with defense, such as flowering time and competitive ability (Agrawal *et al*., 2012), ecological changes in the amount or the type of herbivory may also impact traits beyond herbivory defense. While past studies show that herbicide can reduce the abundance and diversity of herbivores by reducing plant biomass and altering the composition of plant communities (Prosser *et al*., 2016; Capinera, 2018), our study suggests that herbicide treatment of a single plant species may be enough to increase rates of herbivory and alter herbivory patterns, which could lead to longer-term impacts on the herbivore community.

### Absence of a trade-off between herbicide and herbivory resistance

Contrary to our initial expectation, we did not find a trade-off between resistance to glyphosate and resistance to herbivory and instead found they were positively correlated. This result was somewhat unexpected, given that previous work in *Amaranthus hybridus* found a trade-off between resistance to triazine and to herbivory (Gassmann & Futuyma, 2005). However, there are key differences between our study system and the *A. hybridus*/triazine system—the herbicide, plant species, and mechanisms of resistance each differ—such that any of these could be responsible for the different result. In *A. hybridus*, triazine resistance is conferred *via* a target-site mutation in the chloroplast genome that pleiotropically reduces the rate of photosynthesis as compared to susceptible plants, which may consequently weaken plant defenses against herbivores (Gassmann, 2005). Comparatively, *Ipomoea purpurea* exhibits non-target-site resistance to glyphosate, which is most likely conferred through alterations to loci responsible for detoxification (*i*.*e*., cytochrome P450s, glycosyltransferases, and ABC transporters and phosphate transporters; Gupta *et al*., 2023). It is possible that pleiotropy underlies the positive relationship between herbicide and herbivory resistance in *I. purpurea*: in particular, uridine-dependent diphosphate glycosyltransferases (UGTs) mediate the glycosylation of phytohormones integral to herbivory defense (*i*.*e*., abscisic acid, jasmonates, and salicylic acid; (Gharabli *et al*., 2023)) and contribute to herbicide resistance in multiple plant species—including *I. purpurea* (Gupta *et al*., 2023)—through detoxification pathways (Dimunová *et al*., 2022). Additionally, UGTs and cytochrome P450s have been found to be co-expressed in Solanaceae species as part of the biosynthesis of steroidal glycoalkaloids, which contribute to plant defense against herbivory and pathogens (Cárdenas *et al*., 2015). Alternatively, if the likely mechanism of glyphosate resistance in *I. purpurea*—detoxification and altered transport—allows for the shikimate pathway to continue to function in glyphosate-resistant plants, thereby also allowing for the downstream biosynthesis of anti-herbivory phytohormones and defense compounds (Fuchs *et al*., 2021; Gupta *et al*., 2023), the relationship between the two resistance traits may be epistatic rather than pleiotropic. Further functional work discovering the genetic basis of herbivory resistance in this species will be necessary to determine if the positive relationship between the two traits is due to a shared genetic basis or an epistatic relationship between the resistance mechanisms.

### The evolutionary potential of glyphosate and herbivory resistance

Given the absence of a trade-off between the two resistance traits, and yet the presence of a positive correlation between them, one would predict that selection for increased herbicide resistance could lead to higher levels of herbivory resistance, and *vice versa*. We detected the presence of genetic variation for glyphosate resistance—in line with past studies (Baucom & Mauricio, 2008; Debban *et al*., 2015)— suggesting that glyphosate resistance can further evolve if selection is present. We did not, however, find evidence for genetic variation in herbivory resistance in this study population, which is contrary to previous work involving *I. purpurea* that have detected genetic variation for this trait (Rausher & Simms, 1989; Fineblum & Rausher, 1995; Tiffin & Rausher, 1999). Nonetheless, the presence of a significant maternal line by treatment interaction for herbivory resistance in our study population indicates that there is potential for the trait to evolve in the presence of glyphosate, and most likely reflects the genetic correlation between herbivory and herbicide resistance.

In addition to the necessity of genetic variation for the evolution of resistance, the potential for increased herbicide or herbivory resistance is contingent on a link with fitness. Through structural equation modeling, we uncovered a significant relationship between glyphosate resistance and the likelihood of setting seed, but did not find evidence for such a relationship between herbivory resistance and fitness. Thus, while the best-fitting SEM suggests that lineages that are more resistant to glyphosate are more fit, we found no evidence of a fitness benefit of herbivory in this study. We further found that the model with the lowest AIC value included a causal rather than correlative relationship between the two forms of resistances such that higher levels of glyphosate resistance should lead to higher levels of herbivory resistance over time.

Previous work has shown positive selection on glyphosate resistance in field settings in this species (Baucom & Mauricio, 2008), and has shown that glyphosate resistance responds to artificial selection (Debban *et al*., 2015). Our SEM approach in this work furthers our understanding of the relationship between glyphosate resistance and fitness by delineating the direct and indirect effects of glyphosate on the likelihood of seed set in this species. We show that the direct effect of glyphosate (−0.468) is the majority of the total effect of glyphosate application on fitness (−0.672), and is not mediated through foliar damage. Such direct effects may include the truncation of the apical meristem, which stops the plant from growing new vines and delays flowering (Londo *et al*., 2014). The indirect effect of glyphosate on fitness (−0.204) is mediated through foliar damage, as reflected in its negative impact on glyphosate resistance. However, higher levels of glyphosate resistance results in higher fitness, as well as higher levels of herbivory resistance. This suggests that if glyphosate resistance is under selection, higher levels of glyphosate resistance as well as herbivory resistance could evolve through genetic hitchhiking, even if herbivory resistance was not directly under selection.

## Conclusion

The broader aim of our work is to understand how the herbicide glyphosate, a commonly used anthropogenic selective agent, may impact the eco-evolutionary dynamics of plant-herbivore interactions. This study contributes to our broader goals by capturing the ecological impacts of glyphosate on plant herbivory, which could lead to unexpected evolutionary trajectories for both plants and insects. For example, plants may shift their defense strategies to counter increased or different types of feeding following herbicide exposure, whereas herbivores adjust to plants that are physiologically altered by herbicide. Additionally, increases in glyphosate resistance in a population would likewise lead to increased herbivory resistance, which may shift herbivore preferences to less glyphosate-resistant individuals. To better gauge if the ecological shift in insect feeding following herbicide exposure could have evolutionary implications for plant-insect communities, we must better understand the fitness costs and benefits associated with resistance to herbicide and herbivory in plants as well as better understand the subsequent effects of altered feeding on herbivore community composition and behavior.

In conclusion, our study explores how the herbicide glyphosate can alter plant-herbivore interactions and shape the evolution of plant resistance. As with other biotic and abiotic stressors, herbicide has the potential to contribute to the eco-evolutionary dynamics of plant-herbivore interactions by exerting selection on plant defense traits, thereby altering the evolutionary context under which these interactions occur (Iriart *et al*., 2020). Indeed, our study is an example in which herbicide resistance, rather than herbivory resistance, is the likely driver of eco-evolutionary dynamics. It is important to note that different herbicides affect plants through different modes of action, and as such, herbicide resistance is conferred through a variety of different mechanisms (Délye *et al*., 2013; Shaner, 2014). This means anticipating the effects of a particular herbicide on herbivory patterns, or predicting herbivore responses to herbicide resistant plant species could be complicated. By examining the ecological effects of herbicide treatment and the subsequent potential evolutionary implications, we show that a strong anthropogenic stressor, herbicide, has the potential to lead to herbivory resistance, and thus the potential to alter the complex eco-evolutionary feedback between plants and herbivores.

## REFERENCES

Agrawal AA, Gorski PM, Tallamy DW. 1999. Polymorphism in Plant Defense Against Herbivory: Constitutive and Induced Resistance in Cucumis sativus. Journal of chemical ecology 25: 2285–2304.

Agrawal AA, Hastings AP, Johnson MTJ, Maron JL, Salminen J-P. 2012. Insect herbivores drive real-time ecological and evolutionary change in plant populations. Science 338: 113–116.

Agrawal AA, Lau JA, Hambäck PA. 2006. Community heterogeneity and the evolution of interactions between plants and insect herbivores. The Quarterly review of biology 81: 349–376.

Agrawal AA, Maron JL. 2022. Long-term impacts of insect herbivores on plant populations and communities. The Journal of ecology 110: 2800–2811.

Amrhein N, Brigitte Deus, Peter Gehrke, Hans Christian Steinrücken. 1980. The Site of the Inhibition of the Shikimate Pathway by Glyphosate: II. Interference of Glyphosate with Chorismate Formation in vivo and In vitro. Plant physiology 66: 830–834.

Baglan H, Lazzari CR, Guerrieri FJ. 2018. Glyphosate impairs learning in Aedes aegypti mosquito larvae at field-realistic doses. The Journal of experimental biology 221.

Bates D, Mächler M, Bolker B, Walker S. 2015. Fitting Linear Mixed-Effects Models Using lme4. Journal of statistical software 67: 1–48.

Baucom RS. 2019. Evolutionary and ecological insights from herbicide-resistant weeds: what have we learned about plant adaptation, and what is left to uncover? The New phytologist 223: 68–82.

Baucom RS, Mauricio R. 2004. Fitness costs and benefits of novel herbicide tolerance in a noxious weed. Proceedings of the National Academy of Sciences of the United States of America 101: 13386–13390.

Baucom RS, Mauricio R. 2008. Constraints on the evolution of tolerance to herbicide in the common morning glory: resistance and tolerance are mutually exclusive. Evolution; international journal of organic evolution 62: 2842–2854.

Baucom RS, Mauricio R, Chang S-M. 2008. Glyphosate induces transient male sterility in Ipomoea purpurea. Botany 86: 587–594.

Bruce TJA. 2015. Interplay between insects and plants: dynamic and complex interactions that have coevolved over millions of years but act in milliseconds. Journal of experimental botany 66: 455–465.

Capinera JL. 2018. Direct and Indirect Effects of Herbicides on Insects. Weed Control: 76–91.

Cárdenas PD, Sonawane PD, Heinig U, Bocobza SE, Burdman S, Aharoni A. 2015. The bitter side of the nightshades: Genomics drives discovery in Solanaceae steroidal alkaloid metabolism. Phytochemistry 113: 24–32.

Chaney L, Baucom RS. 2014. The costs and benefits of tolerance to competition in ipomoea purpurea, the common morning glory. Evolution; international journal of organic evolution 68: 1698–1709.

D’Alessandro M, Held M, Triponez Y, Turlings TCJ. 2006. The role of indole and other shikimic acid derived maize volatiles in the attraction of two parasitic wasps. Journal of chemical ecology 32: 2733–2748.

Debban CL, Okum S, Pieper KE, Wilson A, Baucom RS. 2015. An examination of fitness costs of glyphosate resistance in the common morning glory, Ipomoea purpurea. Ecology and evolution 5: 5284–5294.

De-la-Cruz IM, Cruz LL, Martínez-García L, Valverde PL, Flores-Ortiz CM, Hernández-Portilla LB, Núñez-Farfán J. 2020. Evolutionary response to herbivory: population differentiation in microsatellite loci, tropane alkaloids and leaf trichome density in Datura stramonium. Arthropod-plant interactions 14: 21–30.

Délye C, Jasieniuk M, Le Corre V. 2013. Deciphering the evolution of herbicide resistance in weeds. Trends in genetics: TIG 29: 649–658.

Dewar AM, Haylock LA, Bean KM, May MJ. 2000. Delayed control of weeds in glyphosate-tolerant sugar beet and the consequences on aphid infestation and yield. Pest management science 56: 345–350.

Dimunová D, Matoušková P, Podlipná R, Boušová I, Skálová L. 2022. The role of UDP-glycosyltransferases in xenobioticresistance. Drug metabolism reviews 54: 282–298.

Ehrlich PR, Raven PH. 1964. Butterflies and Plants: A Study in Coevolution. Evolution; international journal of organic evolution 18: 586–608.

Fineblum WL, Rausher MD. 1995. Tradeoff between resistance and tolerance to herbivore damage in a morning glory. Nature 377: 517–520.

Freitas-Silva L de, Araújo HH, Meireles CS, Silva LC da. 2022. Plant exposure to glyphosate-based herbicides and how this might alter plant physiological and structural processes. Botany 100: 473–480.

Fuchs B, Saikkonen K, Helander M. 2021. Glyphosate-Modulated Biosynthesis Driving Plant Defense and Species Interactions. Trends in plant science 26: 312–323.

Gassmann AJ. 2005. Resistance to herbicide and susceptibility to herbivores: environmental variation in the magnitude of an ecological trade-off. Oecologia: 575–585.

Gassmann AJ, Futuyma DJ. 2005. Consequence of herbivory for the fitness cost of herbicide resistance: photosynthetic variation in the context of plant-herbivore interactions. Journal of evolutionary biology 18: 447–454.

Getman-Pickering ZL, Campbell A, Aflitto N, Grele A, Davis JK, Ugine TA. 2020. LeafByte: A mobile application that measures leaf area and herbivory quickly and accurately. Methods in ecology and evolution / British Ecological Society 11: 215–221.

Gharabli H, Della Gala V, Welner DH. 2023. The function of UDP-glycosyltransferases in plants and their possible use in crop protection. Biotechnology advances 67: 108182.

Gupta S, Harkess A, Soble A, Van Etten M, Leebens-Mack J, Baucom RS. 2023. Interchromosomal linkage disequilibrium and linked fitness cost loci associated with selection for herbicide resistance. The New phytologist 238: 1263–1277.

Hamann E, Blevins C, Franks SJ, Jameel MI, Anderson JT. 2021. Climate change alters plant-herbivore interactions. The New phytologist 229: 1894–1910.

Iriart V, Baucom RS, Ashman T-L. 2020. Herbicides as anthropogenic drivers of eco-evo feedbacks in plant communities at the agro-ecological interface. Molecular ecology.

Johnson NM, Baucom RS. 2022. Dicamba drift alters plant–herbivore interactions at the agro-ecological interface. Ecosphere 13.

Koricheva J, Nykänen H, Gianoli E. 2004. Meta-analysis of trade-offs among plant antiherbivore defenses: are plants jacks-of-all-trades, masters of all? The American naturalist 163: E64–75.

Kuester A, Chang S-M, Baucom RS. 2015. The geographic mosaic of herbicide resistance evolution in the common morning glory, Ipomoea purpurea: Evidence for resistance hotspots and low genetic differentiation across the landscape. Evolutionary applications 8: 821–833.

Kuester A, Fall E, Chang S-M, Baucom RS. 2017. Shifts in outcrossing rates and changes to floral traits are associated with the evolution of herbicide resistance in the common morning glory. Ecology letters 20: 41–49.

Labandeira CC, Currano ED. 2013. The Fossil Record of Plant-Insect Dynamics. Annual review of earth and planetary sciences 41: 287–311.

Labandeira CC, Wilf P, Johnson KR, Marsh F. 2007. Guide to Insect (and Other) Damage Types on Compressed Plant Fossils. Version 3.0. Smithsonian Institution.

Lefcheck JS. 2016. piecewiseSEM : Piecewise structural equation modelling in r for ecology, evolution, and systematics. Methods in ecology and evolution / British Ecological Society 7: 573–579.

Londo JP, McKinney J, Schwartz M, Bollman M, Sagers C, Watrud L. 2014. Sub-lethal glyphosate exposure alters flowering phenology and causes transient male-sterility in Brassica spp. BMC plant biology 14: 70.

Lüdecke D. 2018. Ggeffects: Tidy data frames of marginal effects from regression models. Journal of open source software 3: 772.

Marchiosi R, Lucio Ferrarese M de L, Bonini EA, Fernandes NG, Ferro AP, Ferrarese-Filho O. 2009. Glyphosate-induced metabolic changes in susceptible and glyphosate-resistant soybean (Glycine max L.) roots. Pesticide biochemistry and physiology 93: 28–33.

van der Meijden E, Wijn M, Verkaar HJ. 1988. Defence and Regrowth, Alternative Plant Strategies in the Struggle against Herbivores. Oikos 51: 355–363.

Motta EVS, Mak M, De Jong TK, Powell JE, O’Donnell A, Suhr KJ, Riddington IM, Moran NA. 2020. Oral or Topical Exposure to Glyphosate in Herbicide Formulation Impacts the Gut Microbiota and Survival Rates of Honey Bees. Applied and environmental microbiology 86.

Motta EVS, Raymann K, Moran NA. 2018. Glyphosate perturbs the gut microbiota of honey bees. Proceedings of the National Academy of Sciences of the United States of America 115: 10305–10310.

Myers JH, Sarfraz RM. 2017. Impacts of Insect Herbivores on Plant Populations. Annual review of entomology 62: 207–230.

Orcaray L, Zulet A, Zabalza A, Royuela M. 2012. Impairment of carbon metabolism induced by the herbicide glyphosate. Journal of plant physiology 169: 27–33.

Peterson R. 2021. Finding optimal normalizing transformations via bestNormalize. The R journal 13: 310.

Prosser RS, Anderson JC, Hanson ML, Solomon KR, Sibley PK. 2016. Indirect effects of herbicides on biota in terrestrial edge-of-field habitats: A critical review of the literature. Agriculture, ecosystems & environment 232: 59–72.

Rausher MD, Simms EL. 1989. THE EVOLUTION OF RESISTANCE TO HERBIVORY IN IPOMOEA PURPUREA. I. ATTEMPTS TO DETECT SELECTION. Evolution; international journal of organic evolution 43: 563–572.

R Core Team. 2023. R: A Language and Environment for Statistical Computing.

Santangelo JS, Ness RW, Cohan B, Fitzpatrick CR, Innes SG, Koch S, Miles LS, Munim S, Peres-Neto PR, Prashad C, et al. 2022. Global urban environmental change drives adaptation in white clover. Science 375: 1275–1281.

Schluter D. 1988. ESTIMATING THE FORM OF NATURAL SELECTION ON A QUANTITATIVE TRAIT. Evolution; international journal of organic evolution 42: 849–861.

Schneider MI, Sanchez N, Pineda S, Chi H, Ronco A. 2009. Impact of glyphosate on the development, fertility and demography of Chrysoperla externa (Neuroptera: Chrysopidae): ecological approach. Chemosphere 76: 1451–1455.

Shaner DL. 2014. Lessons Learned From the History of Herbicide Resistance. Weed Science 62: 427–431.

Simms EL, Rausher MD. 1987. Costs and Benefits of Plant Resistance to Herbivory. The American naturalist 130: 570–581.

Simms EL, Rausher MD. 1989. THE EVOLUTION OF RESISTANCE TO HERBIVORY IN IPOMOEA PURPUREA. II. NATURAL SELECTION BY INSECTS AND COSTS OF RESISTANCE. Evolution; international journal of organic evolution 43: 573–585.

Smith DFQ, Camacho E, Thakur R, Barron AJ, Dong Y, Dimopoulos G, Broderick NA, Casadevall A. 2021. Glyphosate inhibits melanization and increases susceptibility to infection in insects. PLoS biology 19: e3001182.

Tiffin P, Rausher MD. 1999. Genetic Constraints and Selection Acting on Tolerance to Herbivory in the Common Morning Glory. Am. Nat. 154: 700–716.

Turcotte MM, Araki H, Karp DS, Poveda K, Whitehead SR. 2017. The eco-evolutionary impacts of domestication and agricultural practices on wild species. Philosophical transactions of the Royal Society of London. Series B, Biological sciences 372.

Van Etten ML, Kuester A, Chang S-M, Baucom RS. 2016. Fitness costs of herbicide resistance across natural populations of the common morning glory, Ipomoea purpurea. Evolution; international journal of organic evolution 70: 2199–2210.

Vila-Aiub MM, Neve P, Powles SB. 2009. Fitness costs associated with evolved herbicide resistance alleles in plants. The New phytologist 184: 751–767.

Wu JC, Xu JX, Liu JL, Yuan SZ, Cheng JA, Heong KL. 2001. Effects of herbicides on rice resistance and on multiplication and feeding of brown planthopper (BPH), Nilaparvata lugens (StÅl) (Homoptera: Delphacidae). International Journal of Pest Management 47: 153–159.

Xie M, Zhang J, Tschaplinski TJ, Tuskan GA, Chen J-G, Muchero W. 2018. Regulation of Lignin Biosynthesis and Its Role in Growth-Defense Tradeoffs. Frontiers in plant science 9: 1427.

