## Supplementary information for "Herbicidal interference: glyphosate drives both the ecology and evolution of plant-herbivore interactions"

**SUPPLEMENTARY TABLES AND FIGURES**

**Table S1.** Linear mixed model for overall herbivory damage, with treatment, and leaf count at the time of glyphosate application (as a proxy for plant size) included as fixed effects. Maternal line by treatment interaction, maternal line, and block were included as random effects, and evaluated using the χ^2^ difference test. Bolded p-values denote statistical significance.

| herbivory damage ~ (treatment × 1\|maternal line) + treatment + leaf count + 1\|maternal line + 1\|block | | | |
| --- | --- | --- | --- |
| **fixed effect** | **df** | **F** | **p** |
| treatment | 1 | 144.26 | **<0.001** |
| leaf count | 1 | 0.14 | 0.517 |
| **random effect** | **df** | **𝟀2** | **p** |
| maternal line × treatment | 2 | 132.45 | **<0.001** |
| maternal line | 1 | 0 | 1.000 |
| block | 1 | 14.84 | **<0.001** |

**Table S2.** Linear mixed models for the effects of glyphosate application, glyphosate resistance, and genetic variation on occurrences of chewing damage types (*i.e.,* hole-, margin-, and surface-feeding). Since we scored each leaf photographed for herbivory data for the presence/absence of each damage type, a leaf could have any and all of the three damage types present. Thus, we are evaluating the *proportion* of damaged leaves with each damage type present rather than the *amount* of each type of damage. The outlined model is a template for these analyses; the fixed effects listed in the table were independently evaluated using the model template, with maternal line and block included as random effects. The exception is genetic variation—there was no fixed effect in this model, and we used the χ^2^ difference test to evaluate the effect of maternal line. Bolded values denote statistical significance.

| damage type ~ effect + 1\|maternal line + 1\|block | | | | | | | | | |
| --- | --- | --- | --- | --- | --- | --- | --- | --- | --- |
| **effect** | **hole-feeding** | | | **margin-feeding** | | | **surface-feeding** | | |
|  | F | df | p | F | df | p | F | df | p |
| glyphosate application (treatment) | 100.49 | 1 | **<0.001** | 346 | 1 | **<0.001** | 5.21 | 1 | **0.023** |
| glyphosate resistance  (1 - damage) | 1.44 | 1 | 0.231 | 1.97 | 1 | 0.162 | 1.46 | 1 | 0.228 |
|  | 𝟀2 | df | p | 𝟀2 | df | p | 𝟀2 | df | p |
| genetic variation  (maternal line) | 0 | 1 | 1 | 0 | 1 | 1 | 0 | 1 | 1 |
| maternal line × treatment | 87.84 | 2 | **<0.001** | 277.75 | 2 | **<0.001** | 0 | 2 | 1 |

**Table S3.** Two types of linear models showing genetic variation for glyphosate resistance. The linear mixed model has glyphosate resistance (1 - *p*) as the response variable, with leaf count (as a proxy for plant size) as a fixed effect, and maternal line and block as random effects. The generalized linear model includes plant death as a binomial response variable (dead = 1, alive = 0), with leaf count as a fixed effect, and maternal line nested in population and block as random effects. All effects were evaluated using the χ^2^ difference test. Bolded p-values denote statistical significance.

| **Linear mixed model** |  |  |  |
| --- | --- | --- | --- |
| glyphosate resistance ~ leaf count + 1\|maternal line + 1\|block | | | |
| **effect** | **df** | **𝟀2** | **p** |
| leaf count | 1 | 2.04 | 0.153 |
| maternal line | 1 | 16.73 | **<0.001** |
| block | 1 | 0 | 1 |
| **Generalized linear mixed model (family: binomial)** | | | |
| death ~ leaf count + 1\|maternal line + 1\|block | | | |
| **effect** | **df** | **𝟀2** | **p** |
| leaf count | 1 | 3.04 | *0.081* |
| maternal line | 1 | 24.06 | **<0.001** |
| block | 1 | 0 | 1 |

**Table S4.** Hypothesized SEMs that were tested as part of model selection. Model A included relationships between all four variables of treatment, herbivory resistance, glyphosate resistance, and plant survival to seed set. Model B omitted the relationship between herbivory resistance and seed set; bolded text denotes this is the model ultimately selected. Model C was modified from Model B such that correlative error was specified between glyphosate resistance and herbivory resistance, assuming both resistance traits are influenced by an underlying process. All three models are reported with AIC scores, as well as 𝟀^2^ and Fisher’s C statistics when applicable.

| **Model** | **Description** | **AIC** | **k** | **n** | **Chisq** | **df** | **P** | **FisherC** | **df** | **P** |
| --- | --- | --- | --- | --- | --- | --- | --- | --- | --- | --- |
| A | Saturated model with all relationships | 4074.785 | 17 | 747 | 0 | 0 | 1 | NA | 0 | NA |
| **B** | **Unsaturated model with no correlative error** | **4072.855** | **16** | **747** | **0.071** | **1** | **0.791** | **0.47** | **2** | **0.79** |
| C | Unsaturated model specifying correlative error between glyphosate resistance and herbivory resistance | 4079.929 | 15 | 747 | 0.071 | 1 | 0.791 | 0.47 | 2 | 0.79 |

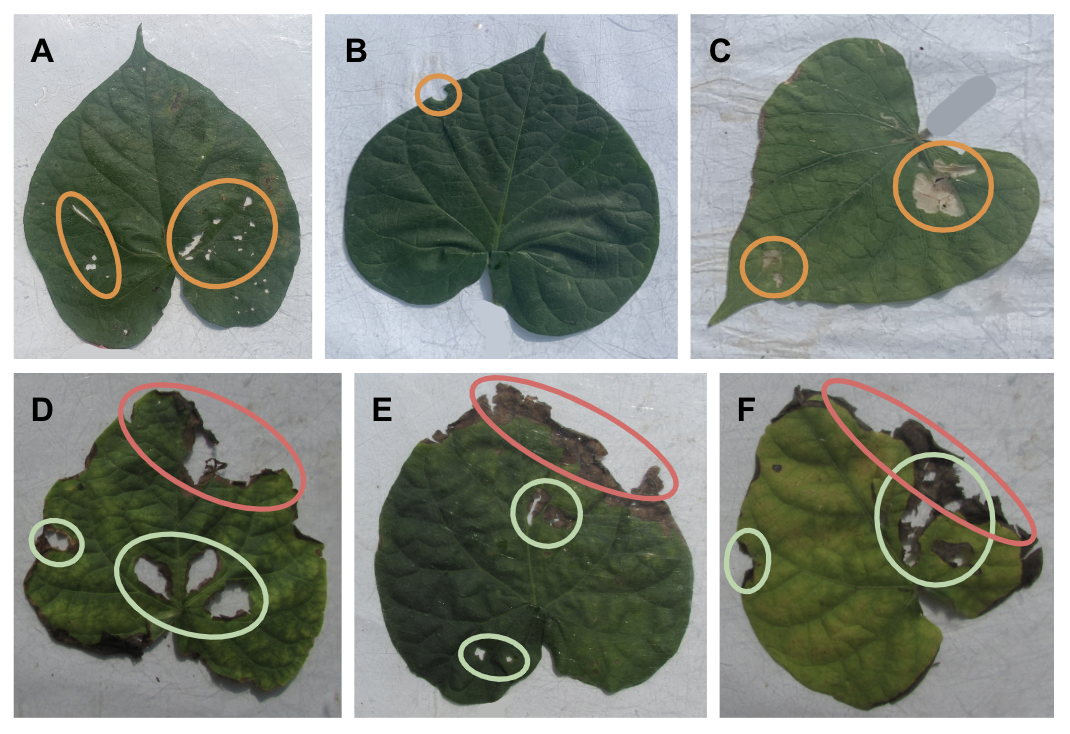

**Figure S1.** Examples of chewing damage types (A) hole-, (B) margin-, and (C) surface-feeding are circled in orange. (D), (E), and (F) are examples of treated leaves missing tissue due to both insect herbivory (green circles) and glyphosate damage (red circles). We took care to differentiate between herbivory and glyphosate damage by excluding missing plant tissue that could not be differentiated between the two causal agents from the herbivory estimate, leading to a more conservative estimation of herbivory damage. To ensure that we were not misattributing glyphosate damage as herbivory damage, we scored each leaf photo from the treatment environment for the presence/absence of ambiguous tissue damage, then built a linear mixed model with this presence/absence as the response variable, glyphosate damage and herbivory damage as fixed effects, and maternal line and block as random effects. We found that glyphosate-susceptible plants had more leaves with the ambiguous damage (F_1_ = 35.113, p < 0.001), while there was no relationship between herbivory damage on each leaf and the presence of ambiguous damage (F_1_ = 1, p = 0.457). This suggests that the presence of ambiguous damage did not interfere with our estimation of herbivory damage and is indeed linked to herbicide damage.

We further validated our result that plants treated with glyphosate experienced higher levels of insect herbivory as compared to untreated plants (Figure 2a, Table S1) by scoring all leaf photos for the presence/absence of herbivory damage, then compared between treatments. We built a generalized linear mixed model with the presence/absence of herbivory as the binomial response variable, treatment as a fixed effect, and block as a random effect, and assessed the model by comparing it to a simplified model without the fixed effect using a χ^2^ test of difference. We found that treatment indeed had an effect on the presence of herbivory (χ^2^_1_ = 150.69, p < 0.001). We then calculated the predicted probabilities of herbivory being present in both treatment environments using the ggeffects package in R (Lüdecke, 2018). We found that leaves treated with glyphosate had a probability of 0.88 of having herbivory damage (95% CI upper limit = 0.93, lower limit = 0.82), while the probability of leaves in the control environment having herbivory damage was 0.67 (95% CI upper limit = 0.77, lower limit = 0.55). This again shows that glyphosate-treated plants indeed experience higher levels of herbivory than plants in the control environment.
